# Insulin resistance, cognition, and functional brain network topology in older adults with obesity

**DOI:** 10.1101/2024.11.26.625552

**Authors:** Clayton C. McIntyre, Robert G. Lyday, Yixin Su, Barbara Nicklas, Sean L. Simpson, Gagan Deep, Shannon L. Macauley, Christina E. Hugenschmidt

**Affiliations:** Neuroscience Graduate Program, Wake Forest Graduate School of Arts and Sciences, Winston-Salem, NC, USA; Department of Radiology, Wake Forest University School of Medicine, Winston-Salem, NC, USA; Department of Cancer Biology, Wake Forest University School of Medicine, Winston-Salem, NC, USA; Department of Gerontology and Geriatric Medicine, Wake Forest University School of Medicine, Winston-Salem, NC, USA; Department of Biostatistics and Data Science, Wake Forest University School of Medicine, Winston-Salem, NC, USA; Department of Physiology and Pharmacology, Wake Forest University School of Medicine, Winston-Salem, NC, USA; Department of Physiology, University of Kentucky, Lexington, KY USA

## Abstract

**Objective:** Cross-sectional data from a sample of older adults with obesity was used to determine how peripheral and neuronal insulin resistance (IR) relate to executive function and functional brain network topology.

**Methods:** Older adults (n=71) with obesity but without type 2 diabetes were included. Peripheral IR was quantified by HOMA2-IR. Neuronal IR was quantified according to a proposed neuron-derived exosome-based method (NDE-IR). An executive function composite score, summed scores to the Auditory Verbal Learning Test (AVLT) trials 1-5, and functional brain networks generated from resting-state functional magnetic resonance imaging were outcomes in analyses. We used general linear models and a novel regression framework for brain network analysis to identify relationships between IR measures and brain-related outcomes.

**Results:** HOMA2-IR, but not NDE-IR, was negatively associated with executive function. Neither IR measure was associated with AVLT score. Peripheral IR was also related to hippocampal network topology in participants who had undergone functional neuroimaging. Neither peripheral nor neuronal IR were significantly related to network topology of the central executive network.

**Conclusions:** Cognitive and functional imaging effects were observed from HOMA2-IR, but not NDE-IR. The hippocampus may be particularly vulnerable to effects of peripheral IR.

## Introduction

Insulin resistance (IR) has long been considered a risk factor for cognitive decline in aging adults. IR was first described in type 2 diabetes, where peripheral tissues (such as muscle and fat) show reduced sensitivity to insulin, resulting in hyperglycemia. Decades of human research has shown that peripheral IR is related to unfavorable aging outcomes [1–4] including lower cognitive performance [5–7], reduced brain glucose uptake [8, 9], and altered functional connectivity in several brain regions, including the hippocampus [10, 11]. These findings suggest that peripheral IR may be involved in brain changes that contribute to cognitive decline. The search for mechanisms explaining the relationship between peripheral IR and brain function has contributed to interest in brain IR, which is the failure of cells located in the brain (including neurons, glia, and vascular cells) to respond to insulin. However, it remains unclear how peripheral IR may contribute to brain IR and cognitive decline [12].

There is histological evidence that Alzheimer’s disease (AD) patients have elevated neuronal IR (defined by dysfunctional phosphorylation of type 1 Insulin Receptor Substrate [IRS-1]), particularly in the hippocampus [13]. However, this finding came from the brains of already-deceased AD patients and therefore cannot be used to study neuronal IR in living humans. To address this limitation, a method for measuring neuronal IR in living humans using neuronal-derived extracellular vesicles (NDEs) was proposed [14]. Data collected using this technique suggest that neuronal IR is elevated in both type 2 diabetes and AD [14]. While these findings suggest that NDEs may be a useful tool for studying neuronal insulin resistance in living humans, outcomes depend upon the surface markers used to isolate NDEs. Currently, there is not a universally accepted surface marker to isolate neuron specific or central nervous system derived extracellular vesicles [15]. Further, only a few published studies have assessed how insulin resistance biomarkers in NDEs (NDE-IR) relate to peripheral IR, cognition, or functional brain connectivity [16, 17].

Insulin receptors are expressed in all cell types in the brain, including neurons of the hippocampus [1, 12, 18, 19]. The hippocampus plays a key role in cognitive aging as it is one of the first brain regions to atrophy in late-onset AD. The hippocampus also expresses the insulin-dependent glucose transport protein GLUT4 at a high density, and findings from the rat brain suggest that insulin signaling through GLUT4 is crucial to cognitive processes including memory formation [20, 21]. Therefore, while it is true that insulin receptors are found in widespread regions of the brain, we hypothesized that effects of neuronal and peripheral insulin resistance may be particularly apparent in the hippocampus.

In the present work, we investigate peripheral and neuronal IR (quantified according to the proposed methods in [14]) as they relate to each other and to brain-related outcomes in a sample of older adults with obesity. First, we assessed whether peripheral and neuronal IR were statistically related with one another. Second, we determined whether peripheral and/or neuronal IR were associated with cognitive abilities. Finally, we determined whether peripheral and/or neuronal IR were associated with functional brain network metrics in regions typically associated with the cognitive outcomes measured in our second aim. Those regions are the central executive network (CEN), which is typically engaged during working memory tasks [22], and the hippocampus, which is well-known for its role in learning and memory formation [23].

## Methods

### Participants

All human subjects research in this study was approved by the Wake Forest University Institutional Review Board. Informed consent was obtained from all participants. All aspects of data collection and analyses in this work adhered to the standards put forth in the Declaration of Helsinki and Belmont reports. All data used for analyses in this work will be made available in de-identified form upon reasonable request from the corresponding author.

Participants were enrollees of INFINITE, a study of weight loss and exercise in older adults (aged 65-80 years) with obesity [24]. Specifically, participants were drawn from an ancillary study, INFINITE MIND, that collected structural and functional brain imaging and cognitive data in a subset of participants [25]. All data included in analyses were from baseline visits of the study, prior to any interventions. Exclusion criteria for the parent study included Mini-Mental State Exam (MMSE) scores < 24, osteoporosis, smoking within the past year, insulin-dependent diabetes, hip fracture, hip or knee replacement, or spinal surgery in the past 6 months, or clinical evidence of depression, heart disease, cancer, liver disease, renal disease, chronic pulmonary disease, uncontrolled hypertension, major physical impairment or contraindication for exercise or weight loss upon exam. Figure 1 demonstrates how participants from the parent study were included in cognitive and functional neuroimaging analyses for this work. Of the 180 participants enrolled in the parent study, NDE-IR was quantified in 85 participants, 14 of which were excluded due to prior diagnosis of type 2 diabetes to avoid confounding effects of diabetes medication. Of the 71 participants without type 2 diabetes, 69 completed cognitive testing and 51 completed structural and functional MRI scans.

**Figure 1.**
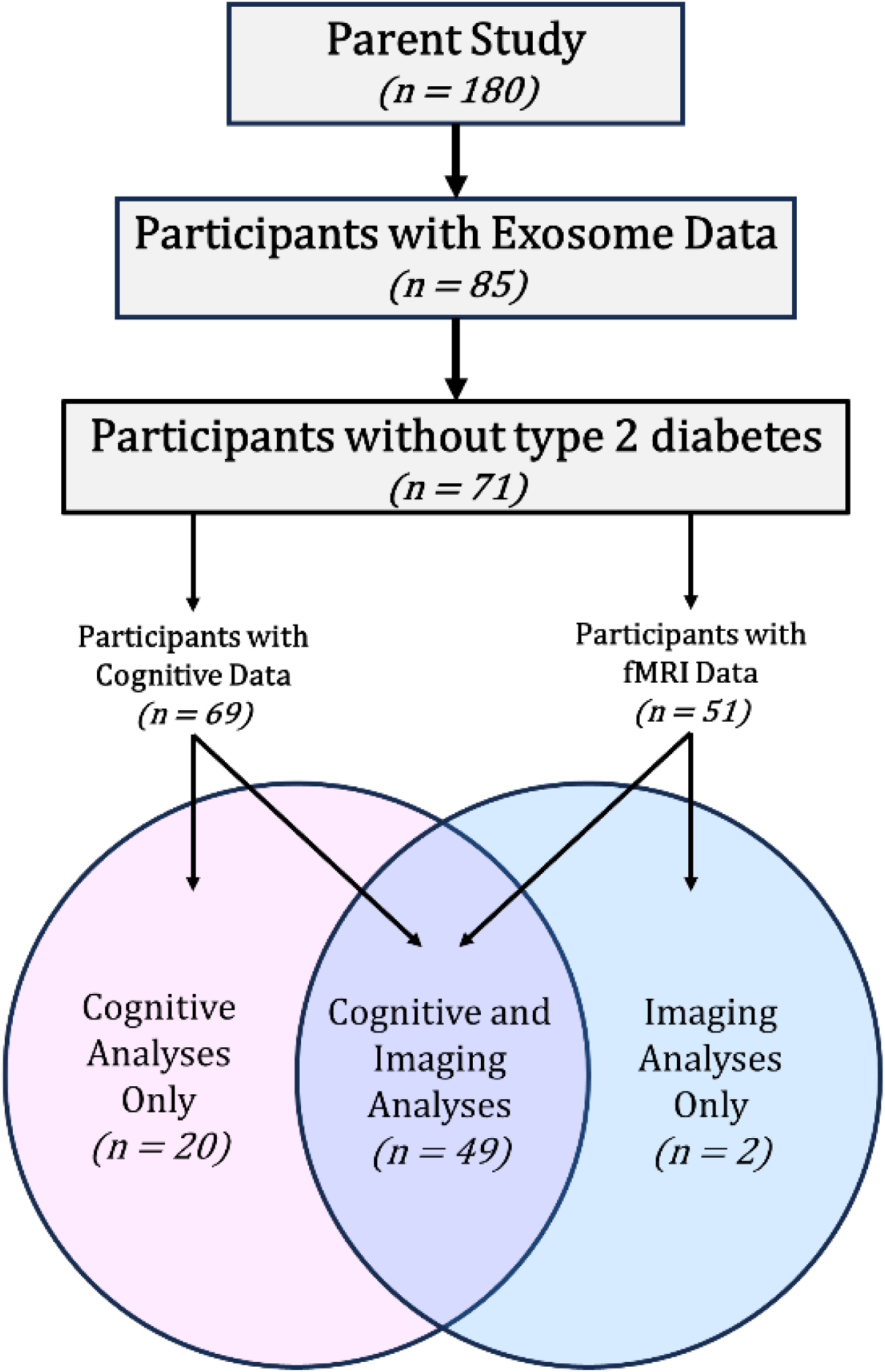
Sample selection for cognitive and imaging analyses. Of the 180 participants enrolled in the parent study, 69 had sufficient data to be included in cognitive analyses and 51 had sufficient data to be included in functional neuroimaging analyses. 49 of the participants between these two analysis groups were the same.

### Peripheral Insulin Resistance

Participants completed a blood draw following a 12-hour fast at their baseline visit for the parent study. From these blood draws, fasting plasma glucose and fasting plasma insulin were used to calculate index values for the computer-based homeostatic model assessment for insulin resistance (HOMA2-IR)[26]. HOMA2-IR is a widely accepted index of insulin resistance in peripheral tissue.

### Neuronal Insulin Resistance

NDEs were isolated from plasma following earlier established protocols [14, 27, 28]. Approximately 500ul of plasma was incubated with 150ul of thromboplastin-D (Fisher Scientific, Inc., Hanover Park, IL) at room temperature for 60 minutes, followed by the addition of 350ul of calcium- and magnesium-free Dulbecco’s balanced salt solution with protease inhibitor cocktail (Roche Applied Sciences, Inc., Indianapolis, IN) and phosphatase inhibitor cocktail (Pierce Halt, Thermo Scientific, Inc., Rockford, IL). After centrifugation at 1500 × g for 20 minutes, supernatant were mixed with 252 μl of ExoQuick precipitation solution (EXOQ; System Biosciences, Inc., Mountain View, CA), and incubated for 1 hour at 4°C. Resultant total extracellular vesicle (TE) suspensions were centrifuged at 1500 × g for 30 minutes at 4°C and each pellet was resuspended in 200 μl of DBS−2 with inhibitor cocktails.

TE suspensions were incubated with 2 μg of mouse anti-human CD171 (L1CAM, transmembrane L1 cell adhesion molecule) biotinylated antibody (clone 5G3, eBioscience, San Diego, CA) for 1 hour at 4°C, then 25 μl of streptavidin-agarose resin (Thermo Scientific, Inc.) plus 50 μl of 3% bovine serum albumin (BSA) incubation for 2 hours at 4°C. After centrifugation at 200 × g for 10 min at 4°C and removal of the supernatant, each pellet was suspended in 50 µl of 0.05 M glycine-HCl (pH 3.0) by vortexing for 10 seconds, incubated at 4°C for 30 minutes, and re-centrifuged at 200×g for 15 minutes at 4°C. Each suspension then received 450ul M-PER mammalian protein extraction reagent (Thermo Fisher Scientific, Incorporated) that had been adjusted to pH 8 with 1 M Tris HCL (pH 8.6) and contained the cocktails of protease and phosphatase inhibitors. These suspensions were incubated at 37°C for 10 minutes with vortex-mixing before storage at −80°C until use in ELISAs.

Protein concentration was determined using the BCA method as per manufacturer’s protocol (Pierce, Rockford IL). The expression of specific proteins in NDEs were quantified by ELISA kits for human P-serine 312-IRS-1 (Life Technologies Corporation, Carlsbad, CA, USA), human P-pan-tyrosine-IRS-1 (Cell Signaling Technology, Danvers, MA, USA), human total IRS-1 (AMSBIO, Incorporated, Cambridge, MA, USA), according to suppliers’ directions.

An insulin resistance index was calculated based on the methods of [14]. Total IRS-1, pan tyrosine phosphorylated IRS-1, and serine 312 phosphorylated IRS-1 were quantified with ELISA. Values were adjusted by the total amount of protein. The ratio of phosphorylated serine 312 to pan-tyrosine was calculated and used as a neuronal-derived extracellular vesicle insulin resistance index (NDE-IR), a proxy for insulin resistance in neurons of the brain.

### Neuropsychological Testing

Prior to completing the cognitive assessment battery, participants were asked whether they had followed their normal daily eating and medication regimens and completed a finger stick glucose test to confirm blood glucose levels were >60 mg/dl. All participants had glucose levels higher than this cutoff.

An executive function composite score was calculated based on performance in a cognitive battery including the Digit Symbol Coding task (DSC), the Trail Making Test (TMT) parts A and B, the Stroop task, phonemic fluency, and semantic fluency. The 90-second version of the DSC was administered; the outcome was the number of correct responses. The TMT outcome was difference in time in parts A and B in seconds. The interference score (interference score = [(time(s) needed for subtask 3) − (time(s) needed for subtasks 1+2)]/2) from the 40-item version of the Stroop task was used. For phonemic fluency, participants verbally generated as many words beginning with the letter F as they could in one minute, and then did likewise for the letters A and S. The sum of all 3 trials was used for analysis. For semantic fluency, participants verbally listed as many animals as possible within one minute and then did likewise for kitchen items. The sum of both trials was used for analysis. The composite score was calculated by summing the z-scores of DSC score, TMT B-A time, Stroop interference score, phonemic fluency, and semantic fluency. The sign of the z-score was reversed prior to summing when necessary, so that positive z-scores indicate better performance. The executive function composite was the pre-specified outcome of the INFINITE MIND ancillary.

In addition to the executive function composite score, participants completed the Auditory Verbal Learning Task (AVLT) trials 1-5 to quantify learning ability. The summed score of the first five trials was used for analyses [29].

### MRI Acquisition

Fifty-one participants completed 60-minute MRI scans at baseline. MRI scans were acquired on a 3.0T Siemens Skyra scanner with a 32-channel head coil (Siemens, Germany). High resolution T1-weighted images were acquired using an MPRAGE-GRAPPA sequence (TR = 1900 ms, TE = 2.93 ms, TI = 900 ms, flip angle = 9 degrees, 176 slices). Resting state blood-oxygen level dependent (BOLD) functional magnetic resonance imaging (fMRI) images were acquired using a whole-brain gradient-echo echo planar imaging sequence (35 contiguous slices; slice thickness = 3.75 mm; in-plane resolution = 3.75 × 3.75 mm, TR = 2.0).

### fMRI Preprocessing

T1-weighted images were warped to the Montreal Neurological Institute (MNI) template using the longitudinal image processing pipeline in Advanced Normalization Tools (ANTs) software (ANTs, https://www.nitrc.org/projects/ants). Because this was an interventional study with two timepoints, for each participant, a single-subject (halfway-space) image was created by warping T1 images from timepoint 1 and timepoint 2 together. Single-subject templates were then combined to create a study-specific template using the antsMultivariateTemplateConstruction command. Brain extraction, N4 bias field correction, and tissue segmentation were performed using ANTs Cortical Thickness. The warp between the study-specific template and the MNI template was calculated, and individual T1-weighted images in native space were transformed to MNI space by simultaneously applying transforms from native space to the study-specific template and the study-specific template to the MNI template.

Preprocessing of BOLD fMRI images was performed using FEAT in FMRIB’s Software Library (FSL, www.fmrib.ox.ac.uk/fsl). N4 bias field correction in ANTS was applied to BOLD images prior to preprocessing and the first 10 volumes of the BOLD image were removed to eliminate irregularities in signal due to the stabilization of the magnetic field. The function epireg was used to apply fieldmaps for distortion correction and perform realignment and slice timing correction. Data were band-pass filtered (0.009-0.08 Hz) to further remove physiological noise and low frequency drift from the functional signal. Images were smoothed with a 5mm Gaussian kernel. Potential confounding for non-neural signal due to motion and other physiological signals was addressed by removing global, white matter, and cerebrospinal fluid signal using the compcorr method [30] implemented through the Conn Toolbox [31]. Fluctuations in signal remaining after controlling for head motion were identified using ART software [31] and time points with meaningful deviations in signal were regressed out. BOLD images were coregistered to their native space structural image and then warped to the MNI template using the same parameters used to warp the T1-weighted structural image to MNI space.

### Network Generation and Efficiency Inference

The Pearson correlation coefficient of the BOLD signal time series for every possible pairing of two nodes was calculated and stored in a whole-brain functional connectivity matrix [32]. Values were thresholded and binarized to create an adjacency matrix by keeping only positive edges with predefined edge density *S* = 2.5 for each participant, where *S* = log(*N*)/log(*k*) with *N* being the number of nodes in the network and *k* being average degree. A threshold of S = 2.5 was used because this has previously been shown to produce networks with minimal fragmentation which are comparable to other naturally generated networks [33]. Each participant’s resulting binarized adjacency matrix was used for functional brain network analyses.

Global and local efficiency were computed for each network. Global efficiency can be measured for any node by finding the average inverse of the path length between that node and every other node in the network. Network global efficiency can be calculated by averaging all nodal global efficiency values. Local efficiency for each node is calculated as network global efficiency calculated on the subgraph of nodes which are directly connected to the node for which local efficiency is being calculated. Local efficiency is closely related to clustering [34] and network resilience. For a full review of network efficiency measures, see [35].

### Non-Imaging Statistical Analyses

To determine whether HOMA2-IR and NDE-IR have a linear relationship, we fit a linear regression model with NDE-IR as the outcome and HOMA2-IR, age, sex, years of education, and BMI as covariates in participants with sufficient data to be included in the model (n=69). Because the absence of a relationship between NDE-IR and HOMA2-IR was considered to be as important as the presence of a relationship for this study, we additionally computed the Bayes factor (using the BayesFactor R package) for the correlation between these two variables to gather supporting evidence in favor of or against the null hypothesis that NDE-IR and HOMA2-IR are not linearly related.

To determine whether peripheral and/or neuronal insulin resistance are associated with executive function, HOMA2-IR and NDE-IR were evaluated for associations with the executive function composite score using a linear regression model. This model also controlled for age, sex, years of education, and BMI.

### Imaging Statistical Analyses

We used BANTOR [36], a novel regression framework for brain network distance metrics to measure the associations of HOMA2-IR and NDE-IR with global and local efficiency. First, to examine the associations between IR measures and global efficiency for the central executive network (CEN) and hippocampus, two separate distance regression models (i.e., one model for the CEN and one for the hippocampus) were constructed with a 3-dimensional global efficiency map serving as the dependent variable. The independent variables for each participant were HOMA2-IR, NDE-IR, age, sex, years of education, and BMI. The distances between the global efficiency maps were calculated with Jaccard distance. The Jaccard distance comes from the Jaccardized Czekanowski similarity index [37](also known as the Ružička index [38]). To convert these similarity indices into distance measures, we calculated the Jaccard distance as 1-Jaccardized Czekanowski index. For each independent variable, the distance was an absolute value difference between participants. These distances were computed between every subject pair to generate a distance matrix for each variable in the model. Then, the independent variable distances were regressed against the global efficiency distance using a linear statistical model with individual-level fixed effects [36]. Because every possible pairing of participants is used to fit this regression model, the goal is not to predict a brain related outcome, but to determine whether there is a significant linear relationship between any included independent variables and brain network measures such as global efficiency. The interpretation of a significant positive β value from this framework is that a larger distance between an independent variable (e.g., HOMA2-IR) in two participants would tend to be accompanied by more distant brain network metrics between the two participants.

This process was repeated with local efficiency maps as the dependent variable. Statistical significance was set at α = 0.05 for all analyses. As neuroimaging analyses were exploratory for future hypothesis generation, we considered type-II error to be at least as impactful as type-I error [39] and report p-values which are unadjusted for multiple comparisons.

## Results

### Participant Characteristics

Participant characteristics for the cognitive analysis and functional neuroimaging samples are shown in Table 1. Of the 51 participants included in imaging analyses, 49 were also represented in the cognitive analysis sample.

**Table 1.** Participant Characteristics.

|  | **Cognitive Analyses**<br>**(n = 69)** | **Imaging Analyses**<br>**(n = 51)** |
| --- | --- | --- |
| Age (years) | 68.5 (3.1) | 68.6 (3.1) |
| Female (%) | 75.4 | 68.6 |
| Education (years) | 14.6 (2.5) | 15.1 (2.3) |
| BMI (kg/m<sup>2</sup>) | 35.6 (3.9) | 35.8 (3.8) |
| Peak VO<sub>2</sub> (mL/kg/min) | 17.7 (3.7) | 17.9 (4.0) |
| Systolic Blood Pressure (mmHg) | 132 (15.5) | 132 (16.3) |
| Diastolic Blood Pressure (mmHg) | 73 (10.4) | 73 (10.8) |
| Body Fat (% of Total Body Weight) | 44.4 (6.0) | 44.6 (6.3) |
| HDL Cholesterol (mg/dL) | 60.1 (16.1) | 60 (15.1) |
| Non-HDL Cholesterol (mg/dL) | 130.5 (42.0) | 125 (38.2) |
| HOMA2-IR | 2.01 (1.05) | 1.99 (1.06) |
| NDE-IR | 6.67 (4.23) | 6.41 (3.77) |
| Executive Function Composite Sum | 0.24 (1.92) | 0.59 (3.04) |
| White Race (%) | 66.7 | 70.6 |

Participants were categorized as having diabetes if they were taking a medication for T2D at the time of their baseline visit. Values are reported as Mean (Standard Deviation) unless noted as a percentage in the variable name. *p*-values are from two- sample T-tests.
BMI = Body Mass Index
VO<sub>2</sub> = Volume of Oxygen Consumption during maximal exertion exercise test
HOMA2-IR = Computer-based Homeostatic Model Assessment for Insulin Resistance
NDE-IR = Neuronal-enriched Extracellular Vesicle Insulin Resistance Index

### Relationship of Peripheral and Neuronal Insulin Resistance

We addressed whether peripheral IR and neuronal IR were related. In a linear regression model (controlling for age, sex, years of education, and BMI), HOMA2-IR and NDE-IR were not significantly related (β = −0.105, SE = 0.483, p = 0.829). The Bayes factor of the correlation between NDE-IR and HOMA2-IR (r = 0.014, p = 0.900) was 0.285 ± 0%, suggesting stronger evidence in favor of the null model (i.e., that NDE-IR and HOMA2-IR are not linearly related) than against the null model.

### Cognitive Outcomes

Associations between measures of IR and cognitive scores are shown in Table 2. A linear regression model controlling for age, sex, years of education, and BMI revealed that HOMA2-IR was negatively associated with executive function (β = −0.798, SE = 0.319, p = 0.015, Figure 2a). NDE-IR, which was also a covariate in the same model, was not associated with executive function (β = 0.005, SE = 0.083, p = 0.954, Figure 2b). Neither HOMA2-IR (β = −1.201, SE = 0.946, p = 0.209) nor NDE-IR (β = 0.109, SE = 0.247, p = 0.660) was significantly associated with AVLT scores in a second regression model with AVLT score as the outcome.

**Figure 2.**
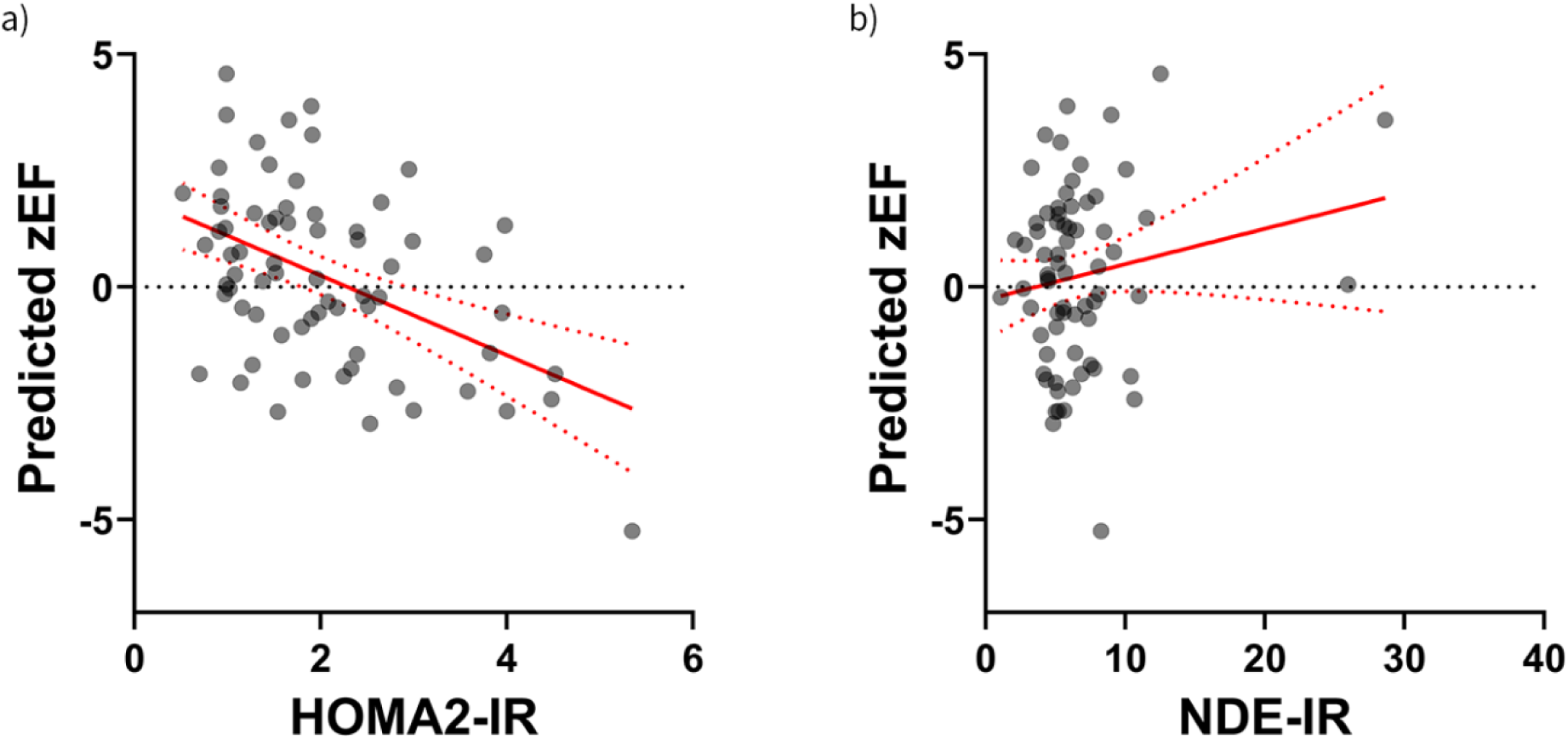
Associations between insulin resistance and executive function. Plots show participants’ insulin resistance levels (x-axes) versus their predicted executive function composite score (y-axis) from linear regression models adjusting for age, years of education, sex, and BMI. Panel **a)** shows the significant negative association relationship between HOMA2-IR and executive function. Panel **b)** shows the statistically insignificant relationship between NDE-IR and executive function.

**Table 2.** IR associations with cognitive measures.

|  | Executive Function Composite |  |  | AVLT Trials 1-5 |  |  |
| --- | --- | --- | --- | --- | --- | --- |
| | $\beta$ | SE | *p* | $\beta$ | SE | *p* |
| Peripheral IR | **-0.798** | **0.319** | **0.015** | -1.201 | 0.946 | 0.209 |
| Neuronal IR | 0.005 | 0.083 | 0.954 | 0.109 | 0.247 | 0.660 |

IR – Insulin Resistance
AVLT – Auditory Verbal Learning Task
SE – Standard Error

### Global and Local Efficiency of Functional Brain Networks

Associations between measures of IR and network efficiency measures are shown in Table 3. HOMA2-IR was not significantly related to global efficiency in the CEN (p = 0.097) or hippocampus (p = 0.724). NDE-IR also was not related to global efficiency in the CEN (p = 0.380) or hippocampus (p = 0.752).

**Table 3.** IR associations with functional brain network topology measures.

|  | Global Efficiency |  |  | Local Efficiency |  |  |
| --- | --- | --- | --- | --- | --- | --- |
| | $\beta$ | SE | *p* | $\beta$ | SE | *p* |
| **Peripheral IR** |  |  |  |  |  |  |
| Central Executive Network | -0.0010 | 0.0006 | 0.097 | -0.0005 | 0.0003 | 0.104 |
| Hippocampus | -0.0004 | 0.0011 | 0.724 | **0.0035** | **0.0014** | **0.011** |
| **Neuronal IR** |  |  |  |  |  |  |
| Central Executive Network | 0.0002 | 0.0003 | 0.380 | <0.0001 | 0.0002 | 0.897 |
| Hippocampus | 0.0002 | 0.0006 | 0.752 | -0.0002 | 0.0007 | 0.782 |

IR – Insulin Resistance
SE – Standard Error

HOMA2-IR exhibited a relationship with local efficiency in the hippocampus (β = 0.0035, SE = 0.0014, p = 0.011). To visualize this result, the average local efficiency maps of participants in the highest and lowest HOMA2-IR tertile are shown in Figure 3. Because HOMA2-IR had been found to be associated with executive function composite scores, a post-hoc BANTOR model with executive function as the outcome and the interaction between HOMA2-IR and hippocampal local efficiency as the independent variable of interest was fit. This model also included age, sex, education, HOMA2-IR, and local efficiency as independent variables. In this model, the interaction between HOMA2-IR and hippocampal local efficiency was significantly related to executive function score (β = 1.943, SE = 0.760, p = 0.011). HOMA2-IR was not significantly related to local efficiency in the CEN (p = 0.104). NDE-IR was not associated with local efficiency in the CEN (p = 0.897) or hippocampus (p = 0.782).

**Figure 3.**
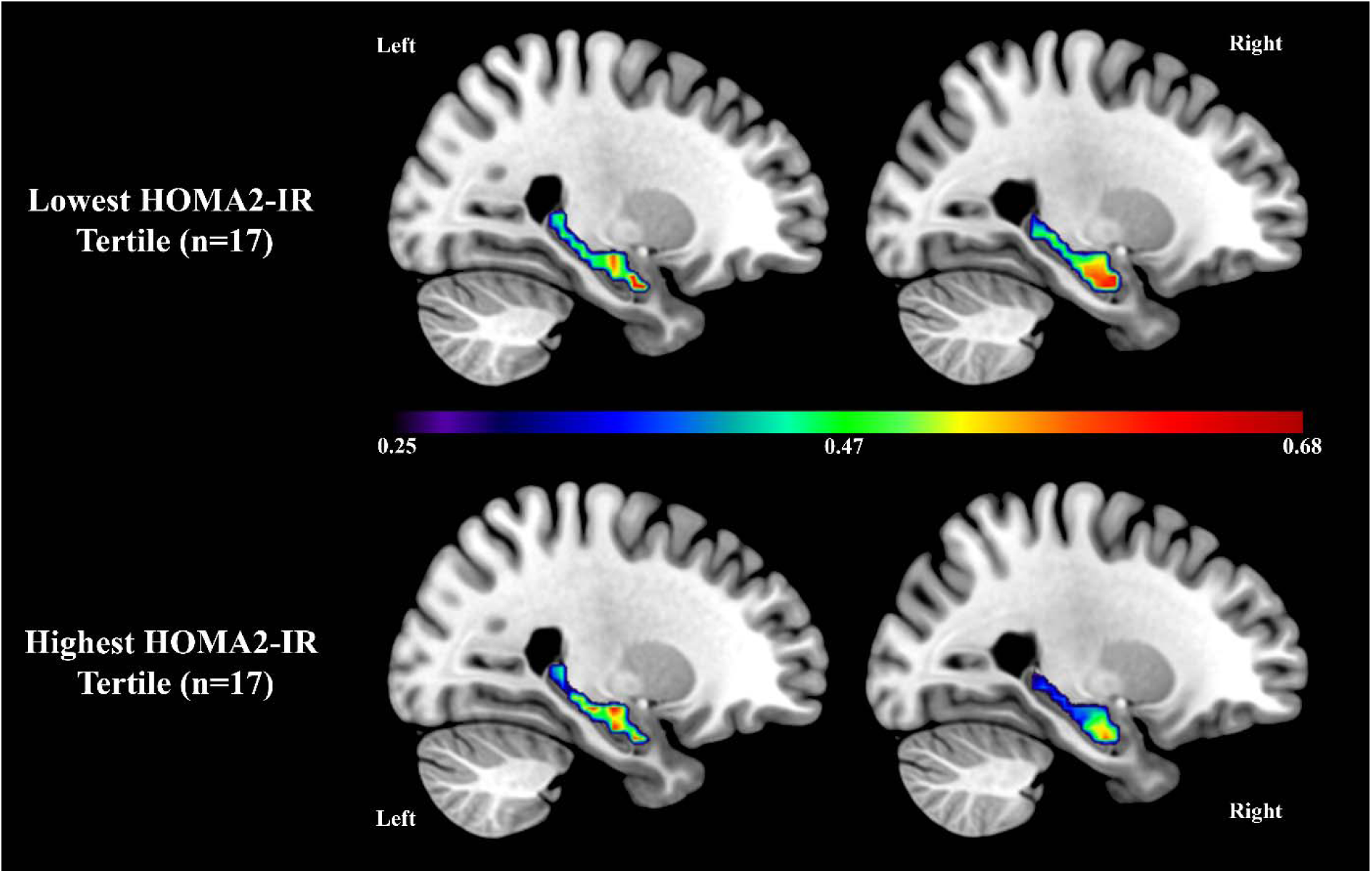
HOMA2-IR and hippocampal local efficiency. Color scale refers to the average local efficiency value of voxels. Warmer colors indicate higher local efficiency. The right hippocampus of participants with low HOMA2-IR has higher local efficiency than in participants with high HOMA2-IR. (MNI Y coordinates: −26 [left], 26 [right])

## Discussion

In this study, we found that HOMA2-IR was not linearly related with NDE-IR (measured using the methods described by [14]) in our sample of older adults with obesity. While we replicated existing findings that peripheral IR is negatively associated with executive function [7, 40] in a unique population, we did not find that NDE-IR was associated with any cognitive outcomes. Finally, functional brain network analyses showed that HOMA2-IR, but not NDE-IR, was related to functional topology of the hippocampus, and the interaction between HOMA2-IR and hippocampal local efficiency was related to executive function ability. We expand on the implications of our findings below.

Previous work [14] demonstrated that NDE-IR is elevated in individuals with type 2 diabetes relative to non-diabetic controls. Data collection for this study occurred soon after publication of the proposed NDE-IR method. In the time between data collection and analyses for the present work, the use of L1CAM as a marker of NDEs has received significant criticism [15]. We chose to include analyses of NDE-IR (which uses L1CAM) in the present work to determine whether this measure was related to any brain-related outcomes of interest in light of its alleged shortcomings. In our study, we did not find any relationship between NDE-IR and HOMA2-IR, cognition, or functional brain network topology. There are caveats to this finding; [14] reported higher NDE-IR values (T2D: 50.4 ± 6.93, non-T2D: 16.4 ± 0.67) compared to ours (non-T2D: 6.47 ± 4.12). Our cohort was four years younger on average than the sample from [14]. A 4-year follow-up visit could show that NDE-IR rapidly increased in the years following the baseline visit in INFINITE. Elucidating the timeframe of NDE-IR escalation longitudinally would be an important next step to inform any future studies of NDE-IR. In addition to the difference in age, the INFINITE sample also features a higher proportion of female participants than [14]. Finally, our study did not have a control group of individuals with a normal BMI or normal HOMA2-IR. It is possible that our cohort was too homogenous for nuanced comparisons to NDE-IR values from other samples.

In functional neuroimaging analyses, we observed a relationship between HOMA2-IR and local efficiency in the hippocampus. Visualizations of this result showed that participants with lower HOMA2-IR had higher local efficiency in their right hippocampus. This finding suggests that an increase in peripheral IR is related to a less functionally-intraconnected hippocampus. In the context of an aging brain, local efficiency offers a quantification of the resiliency of the brain’s functional wiring [35, 41]. More locally efficient networks are expected to minimize the propagation of spurious information through a network. Our finding that differences in HOMA2-IR are related to differences in hippocampal local efficiency may indicate that peripheral IR increases risk for aberrant signaling in the hippocampus. The subsequent finding - that the interaction between HOMA2-IR and hippocampal local efficiency is related to executive function ability - underscores the relevance of peripheral insulin resistance to brain and cognitive function. This finding suggests that intact functional connectivity of the hippocampus may be a protective factor against peripheral IR-related executive function decline. Future work should identify mechanisms through which peripheral insulin resistance impacts hippocampal functional connectivity. Likely mechanisms include oxidative stress and inflammation, though we do not rule out that a novel measure of neuronal insulin resistance may reveal a relationship between neuronal and peripheral IR which was not identified in the present study using an exosome-based approach.

Importantly, we note that the hippocampus has historically been associated with learning and memory formation [23]. The finding that hippocampal connectivity may interact with peripheral insulin resistance to affect executive function (which is typically thought to be governed by the CEN in the frontal and parietal lobes [22]) is at first counterintuitive. Further, peripheral insulin resistance showed no association with scores from AVLT trials 1-5, a well-established measure of learning ability. In interpreting these findings, we note that network neuroscience increasingly suggests that reductive one-to-one mappings of cognitive functions to brain regions (as with learning to the hippocampus) are oversimplifications of more complex, distributed neural processes [42]. For the present study, the implication is that the cognitive impact of changes to hippocampal functional connectivity have the potential to be far-reaching due to the hippocampus’ interdependence with broader subnetworks in the brain. In other words, changes to hippocampal connectivity may impact learning ability, but it may also impact other cognitive domains such as executive function.

Limitations to analyses in this work include the relatively small sample size, the controversial nature of the neuronal IR measure used, the limited assessment of memory ability, and the cross-sectional design of analyses, which prevent us from making conclusions about causality in any of the relationships identified. Notably, we found no evidence that NDE-IR was associated with HOMA2-IR, cognition, or functional brain connectivity. This does not necessarily mean that neuronal IR is not related to cognitive decline. If more reliable measures of neuronal IR are developed, recreating analyses in the present work may provide a more nuanced view of the relationship between neuronal IR and cognitive decline. Additionally, studies could explore the effects of neuronal IR in different populations and assess the impact of neuronal IR on a wider variety of cognitive outcomes.

We conclude by summarizing that peripheral IR was negatively associated with executive function. Additionally, peripheral IR appeared to preferentially alter functional connectivity in the hippocampus, and the statistical interaction between peripheral IR and hippocampal local efficiency was related to executive function ability. In addition to investigation of alternative neuronal IR measures, future work could take advantage of alternative imaging modalities, such as fluorodeoxyglucose positron emission tomography (FDG-PET) imaging or SV2A imaging, which assess glucose uptake and synaptic density, respectively. Ultimately, the findings of this study support the existence of a relationship between peripheral IR and brain function, but limitations to the method for measuring neuronal IR prevent us from making conclusive statements about the role of neuronal IR in cognitive decline.

## Acknowledgements

The authors would like to thank all study participants and their families, as well as study staff for their essential contributions to this work. The dataset analyzed in the current study will be made available from the corresponding author upon reasonable request.

This work was funded by NIH grants R01HL093713 (BJN), K01AG030506 (CEH), UL1TR001420 (Wake Forest Clinical and Translational Science Award), and P30AG21332 (Wake Forest Claude D. Pepper Older Americans Independence Center), American Federation for Aging Research (AFAR) grant RAG12516 (CEH), the Translational Science Center at Wake Forest University, and National Center for Advancing Translational Sciences (NCATS).

## Author Contributions

CEH and BJN provided funding and designed the study. YS and GD carried out lab work needed to quantify exosome-based neuronal insulin resistance from blood samples. RGL and CEH oversaw fMRI data preprocessing. CCM and CEH curated data for analyses. CCM, RGL, SLS, and CEH completed all quantitative analyses. SLM assisted with interpretation of results. CCM wrote the manuscript with input from all authors. All authors contributed to manuscript revision.

